# sORFdb – A database for sORFs, small proteins, and small protein families in bacteria

**DOI:** 10.1101/2024.06.19.599710

**Authors:** Julian M. Hahnfeld, Oliver Schwengers, Lukas Jelonek, Sonja Diedrich, Franz Cemič, Alexander Goesmann

## Abstract

Small proteins with fewer than 100, particularly fewer than 50, amino acids are still largely unexplored. Nonetheless, they represent an essential part of bacteria’s often neglected genetic repertoire. In recent years, the development of ribosome profiling protocols has led to the detection of an increasing number of previously unknown small proteins. Despite this, they are overlooked in many cases by automated genome annotation pipelines, and often, no functional descriptions can be assigned due to a lack of known homologs. To understand and overcome these limitations, the current abundance of small proteins in existing databases was evaluated, and a new dedicated database for small proteins and their potential functions, called ‘sORFdb’, was created. To this end, small proteins were extracted from annotated bacterial genomes in the GenBank database. Subsequently, they were quality-filtered, compared, and complemented with proteins from Swiss-Prot, UniProt, and SmProt to ensure reliable identification and characterization of small proteins. Families of similar small proteins were created using bidirectional best BLAST hits followed by Markov clustering. Analysis of small proteins in public databases revealed that their number is still limited due to historical and technical constraints. Additionally, functional descriptions were often missing despite the presence of potential homologs. As expected, a taxonomic bias was evident in over-represented clinically relevant bacteria.

This new and comprehensive database is accessible via a feature-rich website providing specialized search features for sORFs and small proteins of high quality. Additionally, small protein families with Hidden Markov Models and information on taxonomic distribution and other physicochemical properties are available. In conclusion, the novel small protein database sORFdb is a specialized, taxonomy-independent database that improves the findability and classification of sORFs, small proteins, and their functions in bacteria, thereby supporting their future detection and consistent annotation. All sORFdb data is freely accessible via https://sorfdb.computational.bio.

## 1 Background

A significant portion of bacterial proteins are well studied today, broadly available in public databases, and routinely annotated in newly sequenced genomes [1–3]. Despite these advancements, the exploration of small proteins of up to 100 amino acids (AAs), encoded by short open reading frames (sORFs), has been largely neglected, and they often have been disregarded as noise in eukaryotic and bacterial genomes [1, 4]. Following, we consider small proteins to be functional proteins with a length of 100 AA or fewer.

The application of various length cutoffs for the prediction and identification of protein sequences has led to an inconsistent definition of small proteins. Historically, these cutoffs have resulted from limitations in laboratory protocols and gene prediction tools to reliably detect proteins of such a small size. Gene prediction tools exhibit higher false-positive rates for smaller proteins, which are addressed by implementing strict length limits [5, 6]. Due to the high number of false-positive small proteins in early annotated genomes, minimum length cutoffs were implemented in genome databases [7], and previous small proteins thought to be coding had to be removed later [8].

However, in recent years, the development of experimental ribosome profiling techniques and improvements in mass spectroscopy have resulted in the detection of numerous small proteins [9–11]. Following the identification of small proteins, the elucidation of their purpose has revealed essential cellular functions, including regulatory proteins, membrane-associated or secreted proteins, toxin-antitoxin systems, stress response proteins, and various virulence factors [1, 12–18]. These prominent roles emphasize that the largely unexplored space of small proteins provides essential functions in bacteria. The best-studied bacterial organisms containing small proteins are model organisms and clinically relevant species like *Escherichia coli* and *Salmonella enterica*, in which many new proteins and their encoding sORFs have been reported [10, 18]. The most recently identified proteins in *E. coli* belonged to small proteins with up to 100 AA, particularly with 50 AA or fewer [19].

Consequently, genetic origins and underlying evolutionary mechanisms of small proteins still need to be better understood [17] as they tend to exhibit features differing from genes encoding for proteins longer than 100 AA in bacteria. In particular, start codon usage, ribosomal binding sites (RBSs), and composition biases can differ from longer coding genes [12, 17, 19–21]. These differences could stem from small proteins being developed through *de novo* gene origination, and their comparatively young evolutionary age is insufficient to show the typical organism-specific features of longer coding genes [17, 22]. Pervasive translation of sORFs is also possible [22, 23], although sORFs encoding functional small proteins are probably subject to codon bias [10, 21, 23]. Because of these differences, sORFs and small proteins in bacteria have been overlooked for a long time and are still underrepresented in public databases.

Clustering and identifying new protein families from protein sequences of average length has become a standard bioinformatic procedure. The Markov clustering algorithm [24] has been proven to be reliable for identification in general [25, 26]. However, small proteins challenge existing clustering approaches and tools due to their short length. A metagenomic study has shown vast numbers of hitherto unknown small proteins in human microbiomes and protein families identified by clustering [13].

Addressing these issues, we present sORFdb, to our knowledge the first dedicated database for small proteins and sORF sequences in bacteria. It is a high-quality repository for known sORF and small protein sequences. In addition to protein sequences, physicochemical features are provided to support the search for small protein groups of interest. Furthermore, it offers small protein families and hidden Markov models, enabling the consistent identification and annotation of these families and providing entry points for further research. All data of sORFdb are publicly available for download and can be accessed via an interactive website at https://sorfdb.computational.bio.

## 2 Methods and implementation

### 2.1 Creation of a small protein and sORF database for bacteria

For the creation of the sORFdb database, genomes and protein sequences from various data sources were downloaded and processed. From GenBank (Release 256) [27], the 269,214 latest annotated genomes with an assembly level of “complete genome,” “chromosome,” or “scaffold” were downloaded and used as the primary source for sORFs and small proteins. Small proteins up to 100 AA in length were retrieved from the UniProt database (v2023 03) [2]. Curated and non-fragmented small proteins were downloaded from Swiss-Prot [2], and non-fragmented ones with evidence of existence at the protein, transcript, or homology level were downloaded from UniProtKB [2]. In addition, small proteins of the SmProt database (v2.0) [28] were retrieved and stored with the entries from the UniProt databases in a dataset of verified small proteins. These were directly added to the sORFdb database. For filtering and identification steps of hypothetical proteins and small proteins from an unknown annotation source, the UniRef100 entries [2] of the proteins with evidence and the SmProt proteins were used. Hidden Markov models (HMMs) from AntiFam (v7.0) [8] and Pfam (v35.0) [29] were downloaded, compressed with HMMER (v3.3.2) [30] and used for filtering and scanning for protein domains and motifs.

To extract sORFs and small proteins, several filtering and processing steps were applied to the aforementioned annotated bacterial genomes from GenBank. Complete, unambiguous, and unfragmented sORF and protein sequences and their functional product descriptions were extracted from the annotated genomes. False-positive small proteins were filtered out using PyHMMER (v0.9.0) [31] and AntiFam HMMs [8] with an upper E-value threshold of 1E-5. RBSs of sORFs were detected using Pyrodigal (v2.1.0) [32]. In addition, extracted small proteins were filtered according to whether their annotation source was from a known trusted source, *i.e.*, a reference, representative or NCBI Prokaryotic Genome Annotation Pipeline (PGAP) annotated genome, or an unknown source. Non-hypothetical proteins from trusted annotation sources were stored in the sORFdb database. Additionally, hypothetical proteins and proteins from unknown annotation sources were compared against the dataset of verified small proteins from the Swiss-Prot, UniProt, and SmProt databases using Diamond (v2.1.8) [33].

To examine homology, BLAST Score Ratio Values (SRV), as proposed by Lerat *et al.* [34], were calculated by normalizing bit scores of the best-observed alignment hits with the maximum bit scores of protein self-hits (*Observed score/Maximum score*). This normalization is used because common E-value or bit score thresholds are often too strict for small proteins due to their short length. All homology-filtered small proteins with an SRV of 0.7 or higher were stored in the sORFdb database. To detect very small proteins with only up to 50 AA length, potentially missed by the original genome annotation, a combined approach using Pyrodigal [32], and a homology search with a minimum SRV threshold of 0.7 was employed. A subsequent filtering step excludes all hits overlapping with existing annotations and filters for canonical start codons.

For all small proteins, physicochemical properties were calculated using Biopython (v1.8.1) [35] and Peptides.py (v0.3.1) [36]. Additionally, they were screened for Pfam families and domains with an upper E-value threshold of 1E-5. The taxonomy of all small proteins was adapted to the nomenclature for phyla described by Oren and Garrity [37].

The workflow for the creation of sORFdb was implemented in Nextflow (v23.04.1) [38] to achieve an automated and reproducible procedure. The supplemental materials provide all software packages and tools used, along with their respective versions (Suppl. Tab 1-3).

### 2.2 Clustering of potential small protein families

sORFdb provides potential small protein families along with corresponding HMMs. Due to their short length and understudied clustering properties, a custom graphbased clustering approach was developed to find potential protein families. To reduce the graph size and focus on less studied small proteins, only non-redundant small proteins with 50 AA or fewer were clustered.

During the initial step of the clustering approach, an all-against-all BLAST search was performed using BLAST+ (v2.14.1) [39]. SRVs were calculated for all BLAST hits, and a lower limit of 0.3, as proposed by Lerat *et al.* [34], was applied to identify possible homologs. In addition, a minimum mutual alignment coverage of at least 70.0 % was required to obtain sequence alignments of higher quality and to exclude artifacts of small proteins only sharing a few AA. Afterward, singletons comprising proteins without homologs and distant hits with only one alignment with another protein were excluded. The SRVs of the BLAST hits were transformed into a symmetric undirected graph. For this purpose, SRVs were averaged with their reverse BLAST hit. Small proteins represent nodes and SRVs weighted edges in the graph. To reduce the node degrees in the graph and improve the clustering performance, only *k* best edges of a node were kept. For this purpose, *k* was chosen as the minimum number of edges of a node without creating singletons in the graph. The previous pruning steps exclude distant proteins that otherwise lead to singletons at high values of *k*. Therefore, *k* was chosen as the smallest possible value for which no singletons were reported.

Afterward, to remove edges that could lead to incorrect clusters, a heuristic proposed by Apeltsin *et al.* [40] was applied to every graph component, consisting of a connected subgraph of proteins unconnected to every other connected subgraph.

The pruned graph was then split into batches of components with similar properties depending on the component’s mean node degree and mean edge weights. This was done to improve the selection of the inflation parameter value, which controls the granularity of the clustering. For all batches, a Markov clustering with different inflation values, between 1.2 and 4.0, was computed using MCL (v22-282) [24]. The inflation value was chosen based on the efficiency criterion [41]. A visualization of the clusters used for family identification is available in the supplemental materials (Suppl. Fig. 1).

For all clusters with more than five members, multiple sequence alignments were computed using MUSCLE (v5.1) [42]. Based on these alignments, HMMs were built using PyHMMER (v0.10.2) [31]. Gathering cutoff values were computed for all HMMs, and where possible, a protein product was assigned by a major voting decision based on the annotated protein functions in sORFdb.

### 2.3 sORFdb website

The data of sORFdb is stored in an Elasticsearch cluster [43]. Access to the data is provided via a REST API that was implemented in Java with the Vert.x framework [44]. The website’s graphical user interface was implemented using Vite, Vue, and Typescript [45–47]. It provides a function for an exact sequence and an ID search using the API above to match the queries against the sequences and IDs stored within the Elasticsearch cluster. The alignment-based search uses a BLAST SequenceServer [48] with all stored small proteins as a database returning the IDs of the matching subject sequences which then are used for a search in the Elasticsearch cluster. The small protein family search is performed on the server-side using HMMER (v3.3.2), and the IDs of matching HMMs are also used to search for the HMM entries in the Elasticsearch cluster. For all entries cross-links to the original data sources are provided.

The web-frontend, the Elasticsearch server, and the BLAST SequenceServer are deployed on a scalable Kubernetes cluster, which is hosted in the de.NBI consortium’s cloud computing infrastructure.

## 3 Results

Small proteins encoded by sORFs have long been overlooked in bacteria due to laboratory and computational limitations. With the advent of new laboratory protocols, many small proteins with essential functions have been reported. Despite these advancements and improvements in gene prediction tools, sORFs and small proteins still need to be explored. There are no dedicated databases that focus solely on small proteins in bacteria. sORFdb was created to provide a taxon-independent collection of high-quality bacterial sORFs, small proteins and their assignments to protein families in a comprehensive database to address this issue. This resource is accessible via https://sorfdb.computational.bio.

### 3.1 A large-scale collection of small proteins

To capture bacterial sORFs and small proteins in a comprehensive, taxonomically independent, and standardized manner, including potential unannotated sequences, they were collected from the public data sources GenBank, Swiss-Prot, UniProt, and SmProt, enriched with additional information on taxonomy and RBS usage and processed into a dedicated database. As the public databases contain sequences of varying-quality, all sequences were quality-filtered in a workflow before acceptance.

A total of 31,653,437 sORFs and 34,007,166 small proteins were collected from public databases. 269,214 annotated bacterial genomes from GenBank were systematically screened for sORFs and small proteins, and different filtering steps were applied to extracted sequences. The filtered proteins were split into two groups. The first consists of non-hypothetical small proteins stemming from a trusted annotation source. The second group contains hypothetical ones or ones stemming from an unknown annotation source. From the first group, 22,846,872 annotated small proteins were included in sORFdb. An additional homology-based filtering step was applied to the second group. 8,596,036 small proteins were kept after applying a strict SRV filter since they possessed a homolog with an SRV of 0.7 or higher to sequences from Swiss-Prot, UniProt, or SmProt. For 2,722,346 small proteins previously annotated as “hypothetical protein,” the product description could be updated using information from well-annotated homologs. From UniProt, 2,322,213 small proteins with evidence on transcript, protein, or homology level were collected.

Since automated annotation pipelines rely primarily on computational gene prediction tools, they are limited by hard length cutoffs, and sORFs encoding small proteins may be missed despite possible homologs. To collect these in the annotated genomes, they were detected using a combined approach comprising Pyrodigal, a homolog search, and an overlap and start codon filter. Based on homology alone, 1,363,907 potentially missing small proteins could be detected. After applying the filter mentioned above, a further 198,723 were stored in the sORFdb database. As a result, sORFdb contains 5,073,415 non-redundant small protein sequences and 5,640,450 non-redundant sORF sequences. Despite the absence of sORF sequences for some of the small proteins collected, the total number of non-redundant proteins is smaller than that of non-redundant sORFs due to the use of synonymous codons. Detailed information on the numbers of the total and unique sORF and small protein sequences, as well as related database sources, are shown in Table 1.

**Table 1.**
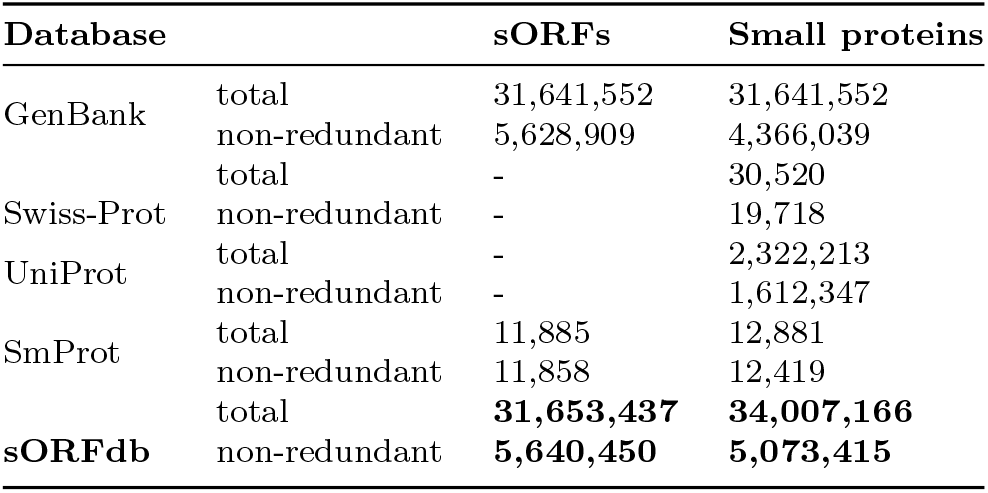
Number of sORFs and small proteins in sORFdb and the used public data sources provides a considerably higher number of non-redundant small proteins, especially with fewer than 50 AA, than the UniRef100 entries with evidence.

| Database |  | sORFs | Small proteins |
| --- | --- | --- | --- |
| GenBank | total | 31,641,552 | 31,641,552 |
|  | non-redundant | 5,628,909 | 4,366,039 |
| Swiss-Prot | total | - | 30,520 |
|  | non-redundant | - | 19,718 |
| UniProt | total | - | 2,322,213 |
|  | non-redundant | - | 1,612,347 |
| SmProt | total | 11,885 | 12,881 |
|  | non-redundant | 11,858 | 12,419 |
| sORFdb | total | **31,653,437** | **34,007,166** |
|  | non-redundant | **5,640,450** | **5,073,415** |

The group of proteins for which the most new proteins have been reported in recent years is the group of small proteins with up to 50 AA [19]. To investigate the length distribution of the total and non-redundant small proteins in sORFdb, their lengths were compared with entries in the UniRef100 database.

In line with expectations, the number of known small proteins in the sORFdb and the UniRef100 database tremendously declines with decreasing length (Fig. 2). The historical and the default gene length cutoffs of standard gene prediction tools and databases (30 AA, 38 AA, and 60 AA) are visible for the predicted UniRef100 entries [5, 7]. In contrast, these cutoffs do neither occur for UniRef100 entries with evidence nor sORFdb entries. While less numerous than all predicted UniRef100 entries, sORFdb

**Fig. 1.**
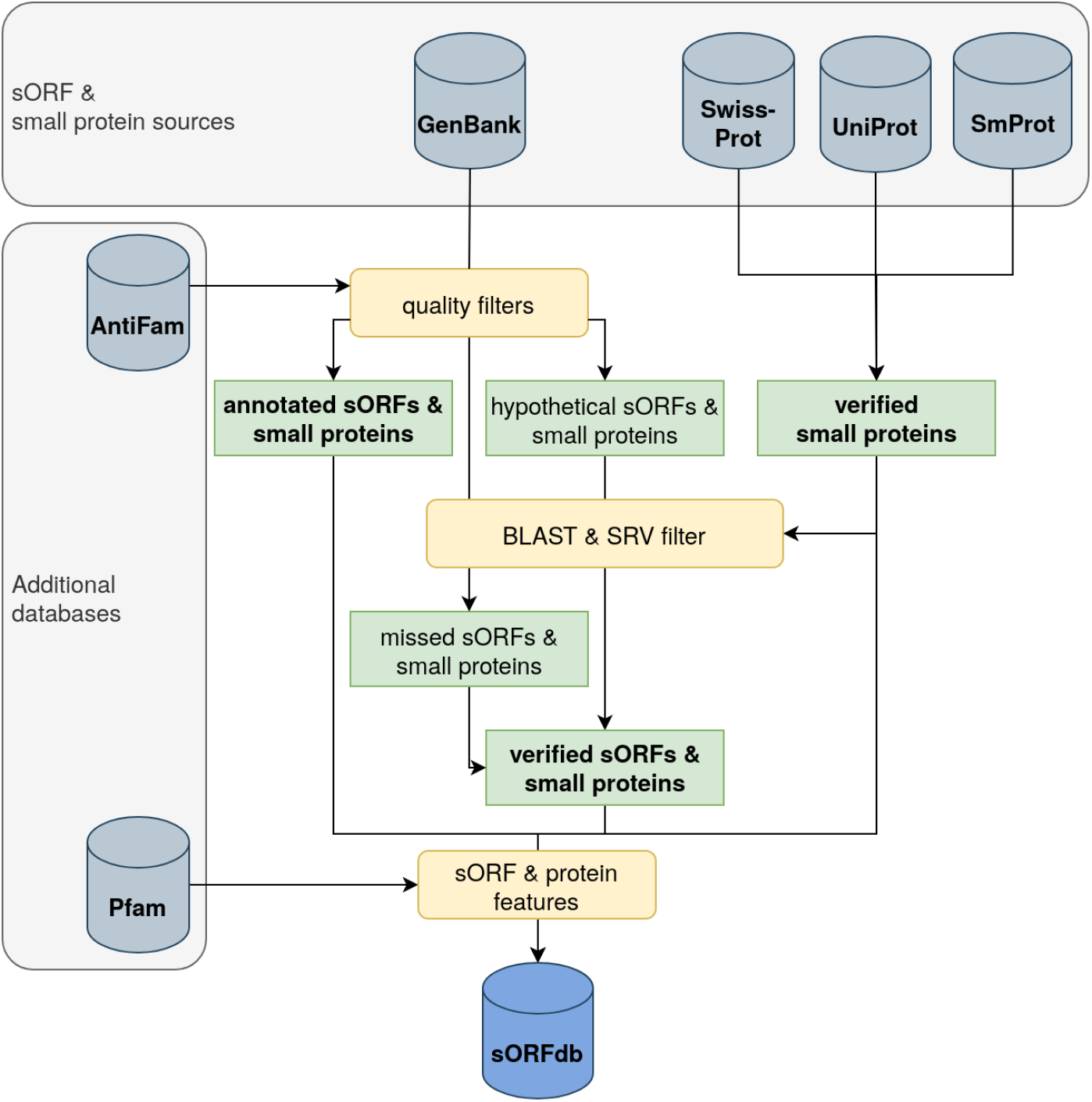
Scheme of the data processing and sORFdb compilation workflow. Annotated genomes from GenBank were quality-filtered for complete and unambiguous sORFs and their annotation source. Small proteins with evidence were retrieved from the Swiss-Prot, UniProt, and SmProt databases and used for sORFdb and additional quality filtering steps. Hypothetical proteins and small proteins from an unknown source were filtered using Score Ratio Value cutoffs based on normalized bit scores. Similarly, missing sORFs were identified. Spurious small proteins were filtered out using AntiFam. Pfam families and domains were assigned, and physicochemical properties were calculated for all small proteins.

**Fig. 2.**
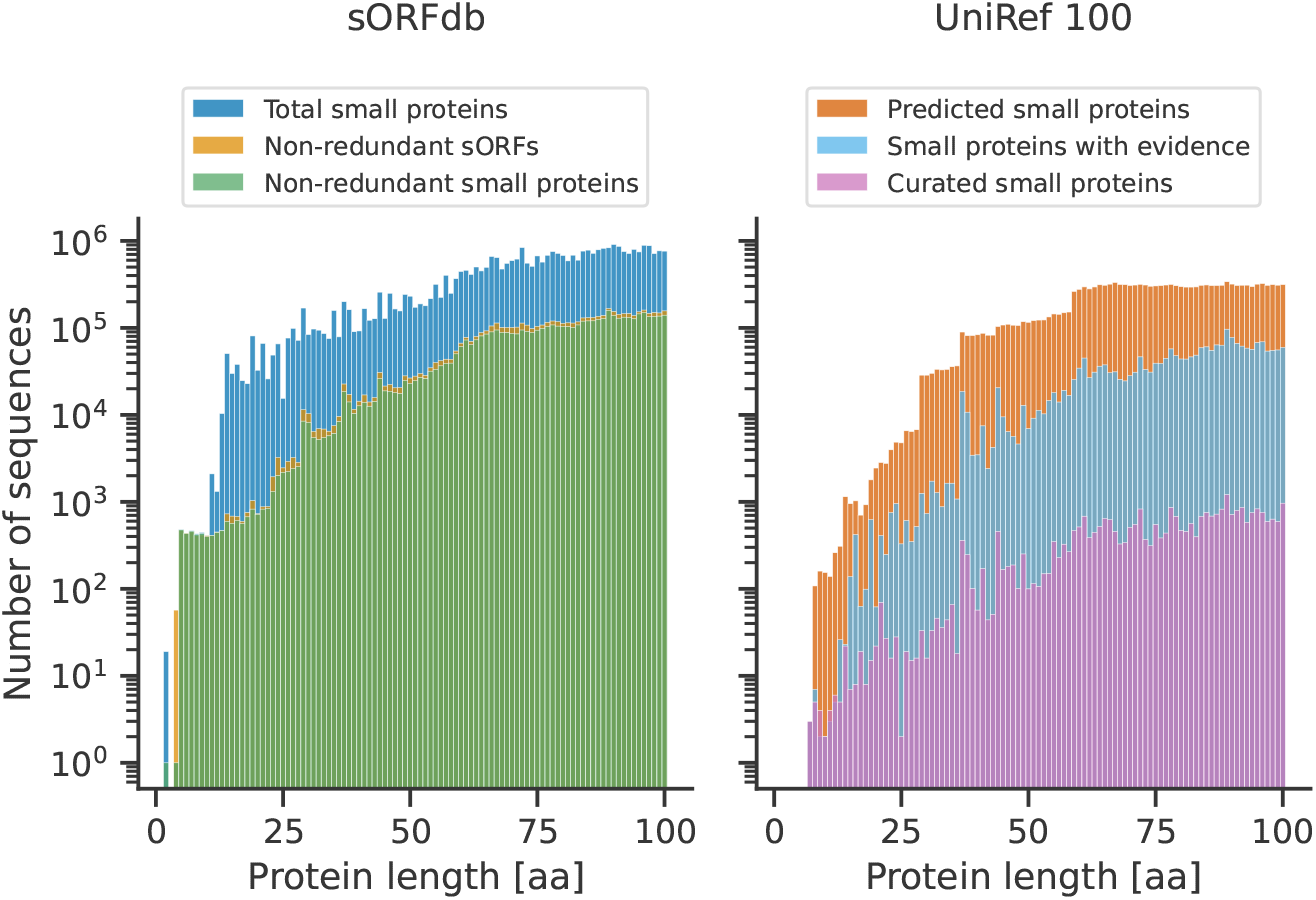
Length distribution of sORFs and small proteins in sORFdb and UniRef100. The number of known small proteins decreases with decreasing sequence length. sORFdb provides more non-redundant small proteins than the UniRef100 database with evidence. Especially for sORFs encoding small proteins with few AA, sORFdb provides more entries.

### 3.2 Taxonomic distribution

Based on the literature and reports for newly identified small proteins in clinically relevant species, we suspected a bias in the taxonomic distribution in our sORFdb database. For this reason, the taxonomy information of sORFs and small proteins was extracted from source genomes and databases to assess the spread and conservation across different bacterial taxa. Phyla were standardized to use a consistent nomenclature based on [37].

In line with our expectation, the taxonomic distribution of small proteins in the sORFdb database showed a clear overrepresentation of clinically relevant species and model organisms (Fig. 3 A). 60.0 % of all small proteins belonged to the phylum of *Pseudomonadota*, formerly known as *Proteobacteria*. Within this phylum, 34.0 % of all protein entries belonged to *Escherichia*, *Klebsiella*, and *Salmonella* genera. Other dominantly represented genera were *Pseudomonas*, *Bacillus*, *Staphylococcus*, and *Streptococcus*, each accounting for 4-6 %. This bias is much less prominent in the taxonomic distribution of non-redundant small proteins (Fig. 3 B). While 42 % of all non-redundant small proteins are also found in *Pseudomonadota*, there are no overrepresented genera, as is the case for all entries in the database.

**Fig. 3.**
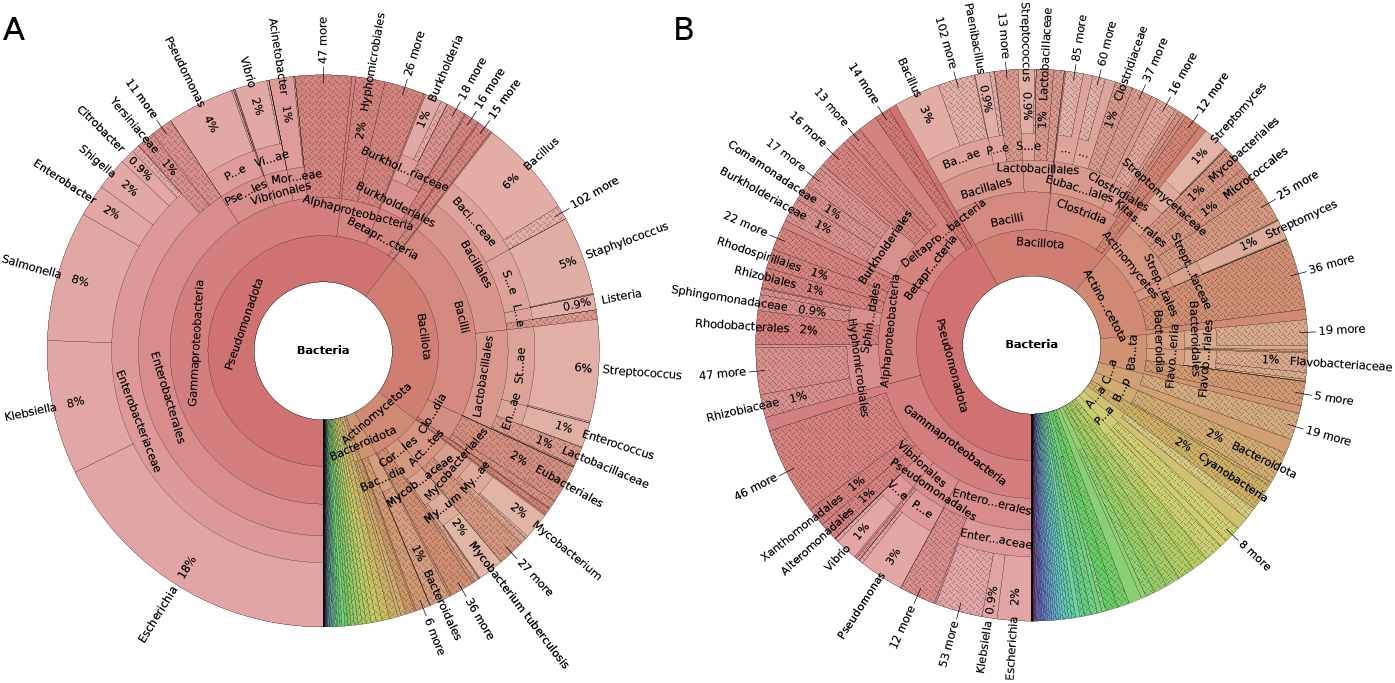
Taxonomic distribution of redundant and non-redundant small proteins. **(A)** Most known small proteins in sORFdb stem from *Pseudomonadota*, model and clinically relevant organisms. **(B)** The taxonomic distribution of non-redundant small proteins showed a much less pronounced bias in comparison. The figure was created with Krona (v2.8.1) [49].

### 3.3 Differing genetic properties between sORFs and longer genes

sORFs are known to have non-canonical start codons more often than genes encoding proteins with more than 100 AA [20]. To determine their start codon usage, these were extracted from all non-redundant sORF sequences in sORFdb. With decreasing length, there was a frequency increase in non-canonical start codons, while canonical start codons were most frequently used for all sORFs encoding small proteins of more than 20 AA (Fig. 4). Although ATG was the most frequent canonical start codon, the frequency of the alternative canonical start codons GTG and TTG increased with decreasing sequence length. sORFs encoding small proteins of 20 AA or fewer had a high proportion of non-canonical start codons compared to longer ones. The codons AAG, ACG, and AGG occur much more frequently in these than in sORFs encoding for small proteins with more than 20 AA. The non-canonical start codons ATA, ATC, ATT, and CTG occurred primarily in these longer sORFs. Regarding their source databases and genera, 73.4 % of the sORFs encoding small proteins with 20 AA or fewer belonged to the genus *Escherichia*. 68.9 % of these sORFs were collected from SmProt, while the remaining sequences were obtained from the GenBank database. In the group with up to 10 AA, 99.8 % of the sequences were annotated in the genus *Escherichia*, and 99.6 % of them were extracted from the SmProt database.

**Fig. 4.**
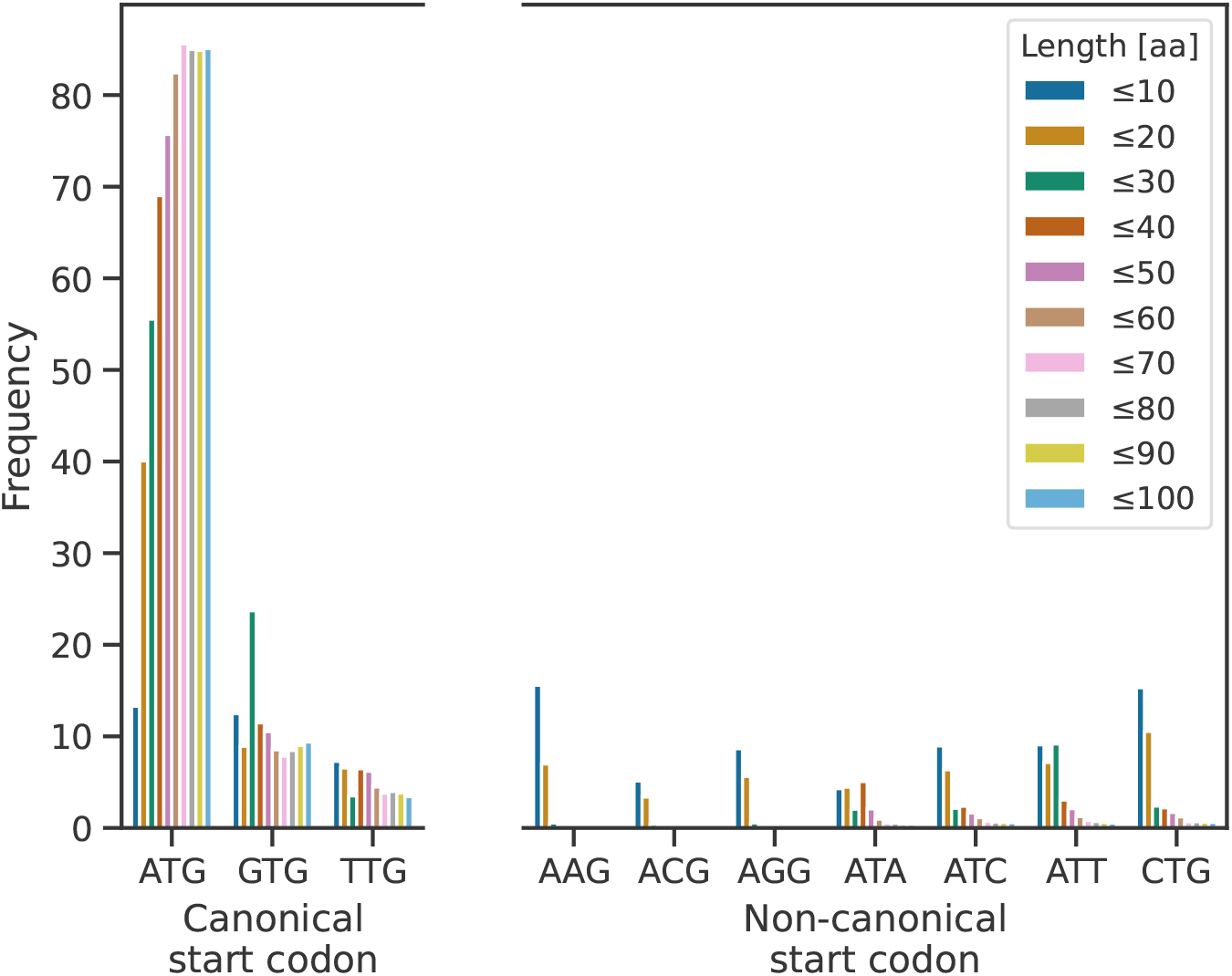
Distribution of canonical and non-canonical start codons in sORFs. With decreasing sORF length, the frequency of non-canonical start codons increases. Shorter sORFs have a higher frequency of the alternative canonical start codons GTG and TTG. The ones encoding small proteins with 20 AA or fewer have the highest frequency of non-canonical start codons. In addition, they show a shift towards different start codons compared to the non-canonical ones used in the group with more than 20 AA.

sORFs are known to be pervasively transcribed, which can happen through leaderless translation [22, 23]. To analyze potential leaderless translation, the usage of RBSs of the non-redundant sORFs in the annotated genomes from GenBank was examined using Pyrodigal [32]. RBS could be detected in 73.8 % of all non-redundant sORFs. Despite this fact, the existence of an RBS varies enormously depending on the sequence length. With decreasing size, the detection of an RBS decreased (Fig. 5). This can be observed for sORFs encoding small proteins with 60 AA or fewer. For all sORFs encoding small proteins with 10 AA or fewer, no RBSs could be detected, and 87.5 % of the ones encoding small proteins with 20 AA or fewer also did not have a predicted RBS.

**Fig. 5.**
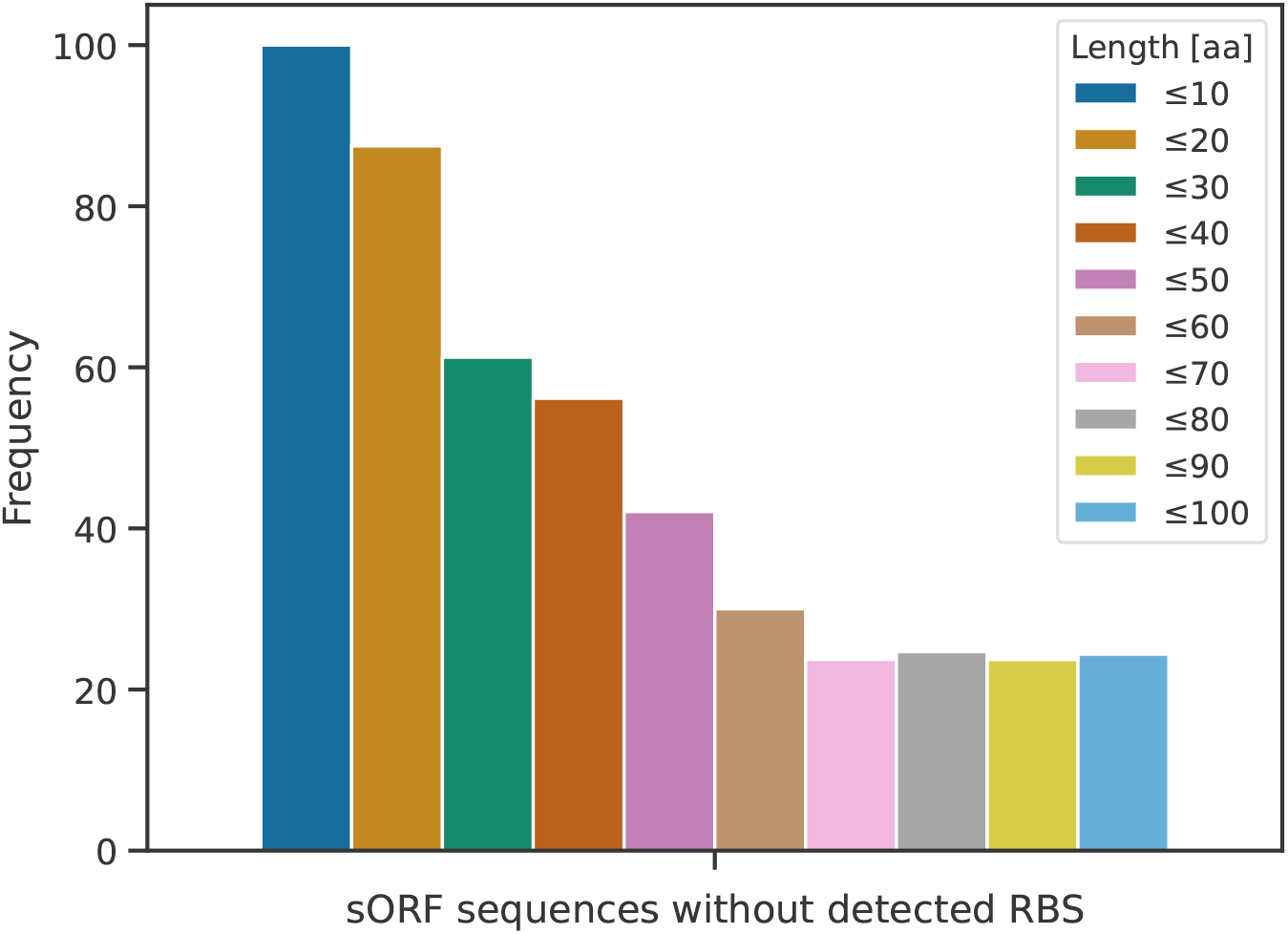
Frequency of sORFs without a detected ribosomal binding site. With decreasing sORF length, the frequency of detected RBSs also decreased, and for sORFs encoding proteins with 10 AA or fewer, no RBSs were found at all.

### 3.4 Functions of small proteins

Functional characterizations of small proteins have revealed essential roles in bacteria. However, homologs and functional descriptions are often unavailable for newly discovered small proteins. For this reason, all small proteins were filtered during the sequence collection process, and hypothetical protein products were re-annotated with functional descriptions of homologs, if available. In addition, all small proteins were queried against the Pfam database to assign protein families and domains.

The most common functional descriptions and Pfam hits were analyzed to investigate whether the functions of the collected small proteins were consistent with the literature. For 74.0 % of the non-redundant small proteins, a Pfam family or domain could be assigned, with the number of known assigned Pfam domains decreasing with decreasing sequence length. Most small proteins in sORFdb are structural proteins of ribosomes that are highly conserved and well-studied. Besides these, essential functions of regulatory proteins, stress response proteins, and toxin-antitoxin systems are predominant. This is in line with previous reported findings [1, 14, 15, 17, 19]. Most regulatory proteins are denoted as helix-turn-helix containing transcriptional regulators. Cold-shock proteins are the most abundant stress response proteins, and the three most common toxin-antitoxin systems are Type II toxin-antitoxin systems of the RelE/ParE, HicA or Phd/YeFM families. The top 20 small protein product annotations are available in the Supplementary Material Table 4. Small proteins with 50 AA or fewer also frequently possess the functional description for helix-turn-helix containing transcriptional regulators. However, many of these are membrane-associated, like the yjcZ family sporulation protein, lmo0937 family protein ATPase subunits, and others. The most common toxin-antitoxin systems are entericidin family proteins. This has also been reported in previous studies [1, 12, 17, 19]. In addition, small proteins with the domain of unknown function DUF3265, DUF2256, or DUF1127 are also frequently found in the annotations of sORFdb and the assigned Pfam domains.

### 3.5 Families of small proteins in bacteria

While small proteins in bacteria are a rapidly evolving field of research and the number and deduced functions of novel identified proteins are the subject of current studies, their families are still understudied [13]. Clusters of similar proteins can be used as a starting point to identify conservation within and across taxonomic groups and the evolution of beneficial functions of these proteins. To address this, potential families were inferred using a custom graph-based clustering approach on the non-redundant collection of bacterial small proteins.

Small proteins with up to 50 AA are the most rapidly growing protein category [19], and longer proteins have been better studied since they were not affected by historical cutoffs [5–7]. For this reason, the clustering and the small protein families focused on the 309,042 less-studied small proteins with up to 50 AA. The clustering approach was developed to handle small protein sequences’ properties better and make minimal assumptions about their clustering behavior. After applying different pruning strategies, 272,018 small proteins remained for the clustering with MCL. 16,518 clusters were assigned in total by the graph-based clustering approach, of which 4,073 were singletons. Clusters with at least five members were used as the basis for the small protein families. Many of the separate graph components were completely assigned to one cluster. There were also large complex-structured graph components, consisting of proteins such as ribosomal or small proteins sharing a domain of unknown function, for which many clusters were determined (Supplementary materials Fig. 1).

Clusters with at least five members were denoted as small protein families to distinguish between technical sequence clusters and potential families. Based on this, 8,884 novel small protein families were created. These families had a mean of 27.7 and a median of 11 members. The most prominent family consisted of 363 members. Most families had members with a length between 40 and 50 AA. Additionally, there were more families with sequences of approximately 38 AA in length. While there was a slightly increased number of families with members of around 30 AA in length, families with shorter members were scarce (Fig. 6). HMMs were built for all small protein families, and accompanying gathering cutoffs were calculated to foster the detection and annotation of small proteins belonging to the identified families. For 8,798 of the 8,884 families, a functional description could be assigned using a majority voting approach based on the existing functional annotation of cluster members. For example, these families shared simple protein motifs, such as a domain of unknown function or a functional description. The most abundant functional descriptions assigned to the families stemmed from ribosomal proteins and regulatory proteins.

**Fig. 6.**
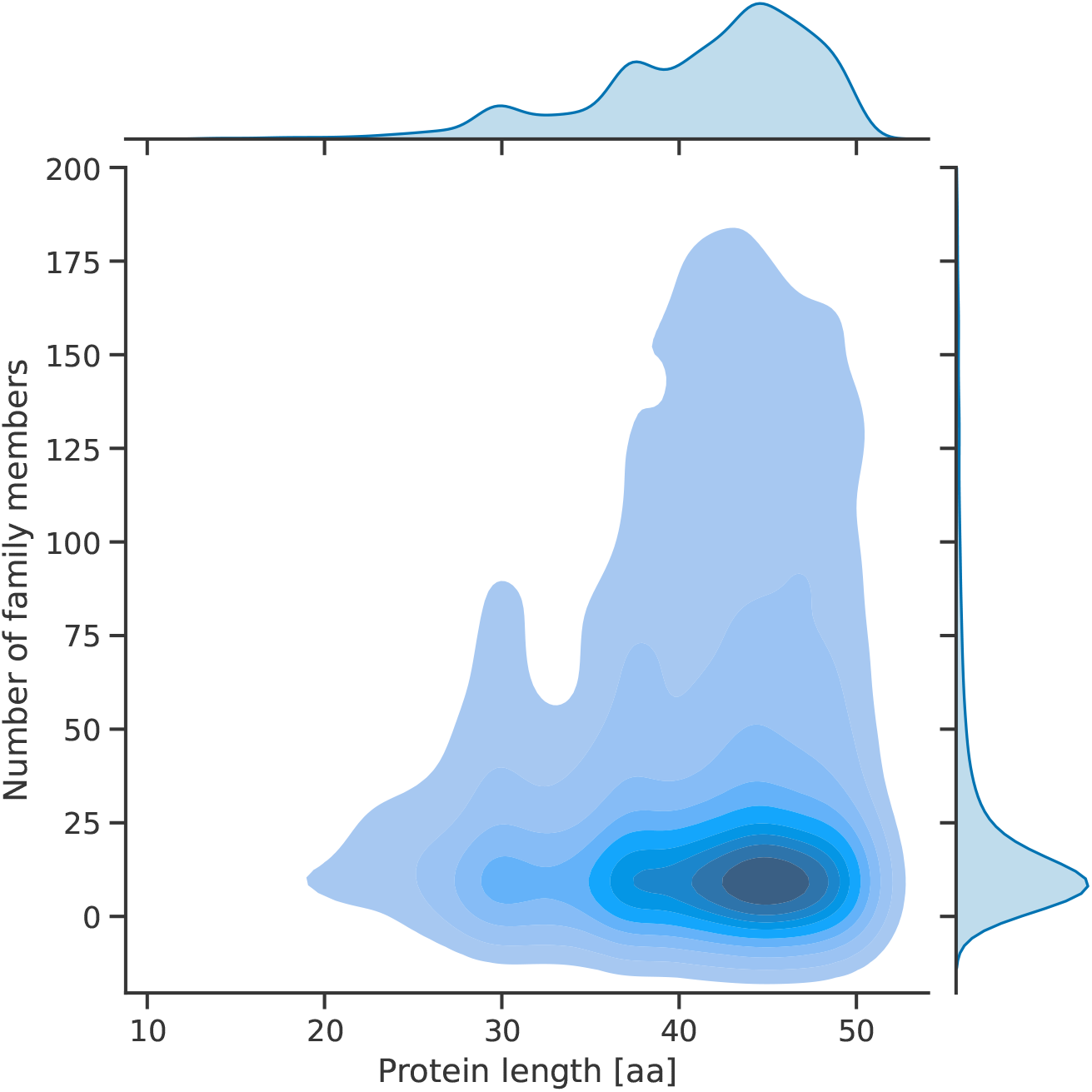
Distribution of cluster size and sequence length of small protein families. Most small protein families had an average member length between 40 and 50 AA or around 30 AA. Most of the families with shorter member proteins had members with a length of about 30 AA.

### 3.6 Interactive web-based access to sORFdb

An interactive website was developed to provide a user-friendly interface for the sORFdb database and to integrate additional services for sequence and family search and data exploration. It makes the collected sORFs, small proteins, small protein families, and related information easily accessible to the scientific community. To accomplish this, it offers various functions for the interaction with the collected data. The database, protein and sORF sequence data, small protein families and the corresponding HMMs are available for download.

To enable researchers to find homologous sORF and small protein sequences, a sequence-based search function is provided with a fast, exact search and a similarity-based BLAST search (Fig. 7 A). In addition, a highly sensitive search for small proteins belonging to known protein families is available (Fig. 7 B). All sequences and families are findable and accessible via unique IDs. Besides sequence-based search functions, browse functions are provided to view small proteins and sORFs matching user selected criteria. These criteria can be based on taxonomy, sequence features, functional description, or physicochemical properties (Fig. 7 C). Search results and filtered sORFdb entries can be downloaded for local processing. To provide further information, links to original resources are provided on a detailed page for each database entry. Similarly, the small protein families can be browsed and inspected.

**Fig. 7.**
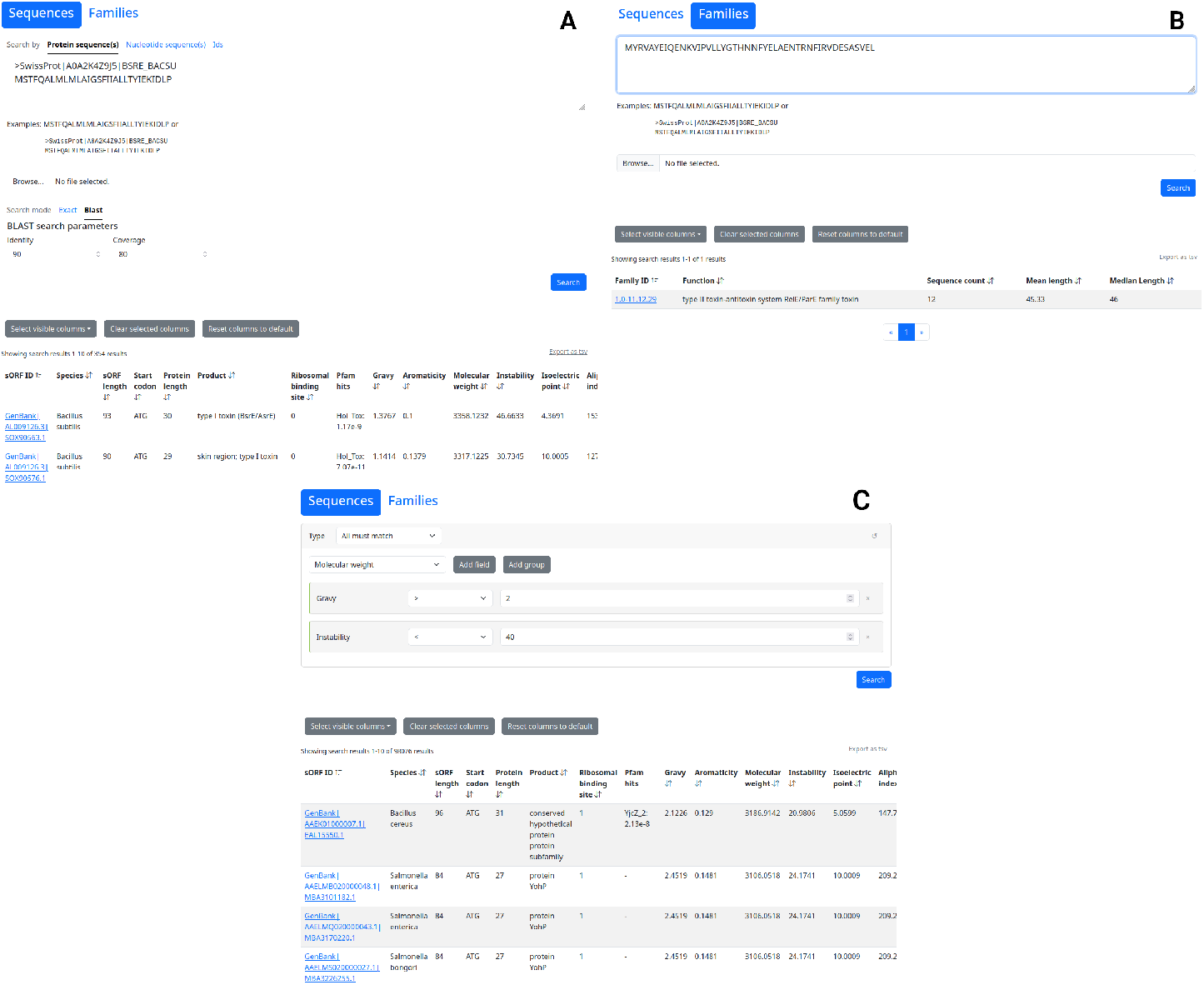
Screenshots of the sORFdb web-frontend (A) The sORFdb website’ search page offers a fast exact and a BLAST-based sequence search for all sORFs and small proteins in the database. **(B)** The small protein family page provides access to the small protein families, their member sequences and properties. **(C)** The browse page provides an interactive selection for taxonomy, sequence-based features, and physicochemical properties to view matching entries in the database.

## 4 Discussion

Small proteins of 100 AA or fewer encoded by sORFs have long been overlooked in bacteria [1, 4]. However, the advent of ribosome profiling, improvements in mass spectrometry, and metagenomics have led to the identification of numerous sORFs and small proteins [9–11, 13]. Despite this, sORFs and small proteins with evidence on transcript or protein level are often missing from public databases and newly annotated genomes. To address these issues, the landscape of publicly available bacterial sORFs and small proteins in annotated genomes and protein databases was captured and analyzed to provide a unified, dedicated database for sORFs, small proteins, and their families.

To the best of our knowledge, sORFdb is currently the largest and most comprehensive sequence database for bacterial sORFs, small proteins, and related families. The combination of different data sources, particularly the integration of GenBank and the application of filtering steps, provides access to a broad collection of sequences and a higher number of sequences than individual data sources used. Due to this approach, sORFdb is taxonomically independent and not focused on specific species compared to smaller databases such as SmProt [28]. In addition to the small protein and encoding sORF sequences, information on RBS usage and physicochemical properties are provided. Most importantly, sORFdb defines families for small bacterial proteins to facilitate the consistent identification of these hard-to-predict proteins as a starting point for further studies. Regarding the functions of small proteins, the annotated functions of the non-redundant proteins in sORFdb largely coincide with the literature. As expected, the most frequently found proteins are ribosomal. The following annotated top functions of regulatory proteins, membrane-associated proteins, stress response proteins and toxin-antitoxin systems of small proteins are also often described in the literature [1, 12–15, 18]. Besides these functions, the Pfam domains DUF3265, DUF2256, and DUF1127 occur with high frequency in small proteins with up to 50 AA. Recent studies show that proteins with the DUF1127 domain serve essential functions concerning the sRNA maturation and RNA turnover as well as the phosphate and carbon metabolism [50, 51]. Based on the DUF1127 small proteins, 37 different families with DUF1127 were identified. In contrast, the 276,091 singletons and 3,561 clusters with up to 4 small proteins identified with the clustering approach show the existence of less conserved or understudied groups. This is also consistent with reports of small proteins conserved in only a few organisms [1]. While the functional annotations of well-studied small proteins in sORFdb align with the known literature, further investigation is needed for less conserved and understudied groups. The taxonomic distribution of the available small proteins in sORFdb shows a clear bias towards Pseudomonadota, particularly towards *E. coli* (Fig. 3 A). This bias is due to the historical over-representation of these bacteria in sequenced genomes and the fact that most ribosome profiling experiments are conducted in this organism [10, 18]. Although this bias is not as evident for the non-redundant small proteins (Fig. 3 B), it still imposes limitations. Combining different databases did not reduce this known bias, and SmProt, which only contains data from *E. coli*, further reinforced this effect. Nevertheless, the taxonomic distribution of non-redundant small proteins shows that sORFdb covers a broad range of non-clinically relevant bacteria despite this bias, allowing a taxon-independent search for small proteins.

Since automated annotation pipelines and gene prediction tools are limited in their ability to predict sORFs, a homology-based approach was used to detect potential missing sORFs in annotated genomes from GenBank. Identifying missing small proteins was based on assumptions derived from the current knowledge of sORFs encoding functional proteins, which is biased towards *E. coli* and related bacteria. Therefore, only potentially missing small proteins with canonical start codons, a known homolog and prediction with Pyrodigal were included [10, 12, 21, 23]. Applying this search filter, 198,723 likely non-spurious small proteins were identified from the 1,363,907 candidates found by homology search. This comparatively low number, in conjunction with the filters applied, and the matching taxonomic bias of the database, indicates that more than a homology search is needed for identifying small proteins. Another limitation is the computational prediction of sORFs since the used tools are not optimized for sequences of such short lengths.

The distribution of start codons in non-redundant sORFs contrasts with the criterion for using canonical start codons that was applied for missing sORFs in the GenBank genomes. As the length decreases, the number of non-canonical start codons increases and shifts towards different non-canonical start codons compared to sORFs that encode small proteins with more than 20 AA (Fig. 4). A possible reason could be that most sORFs of this length were collected from SmProt and identified by ribosome profiling in *E. coli* alone. Therefore, they might include sORFs encoding non-functional transcripts expressed by pervasive translation [21, 22]. Alternatively, these sORFs may be evolutionary young, stemming from *de novo* gene origination, and therefore may not exhibit the typical start codon and RBS usage observed in *E. coli* [17]. To address this, further studies on the codon usage of sORFs encoding functional small proteins are needed to distinguish spurious sORFs expressed by pervasive translation from sORFs stemming from *de novo* gene origination and conserved sORFs.

The selection of an appropriate identity threshold is critical for the clustering of homologous protein sequences. If prior knowledge is available, a suitable threshold can be chosen depending on the evolutionary distance or available information about the composition of the protein families to be clustered. Otherwise, a 30 % identity threshold or the application of E-value or bit score thresholds have been shown to capture more distant homologs [52]. However, this approach cannot be applied to small proteins because for short sequences, even self-hits can have values below these established thresholds [52]. For this reason, a custom graph-based clustering approach was used to identify small protein families. This approach was chosen to minimize assumptions about the clustering behavior of small proteins as much as possible since their clustering properties are not well known. SRVs based on normalized bit scores were used as a similarity metric to address the possibility of insignificant bit scores for shorter small proteins, which can occur even for self-hits. Here, the lower threshold of 0.3 allows the detection of distant homologs. The clustering granularity is automatically selected based on the inflation value with the highest efficiency score. The various pruning steps aim to improve the clustering by excluding singletons and barely matching small proteins beforehand.

A total of 8,884 small protein families with at least five members were identified using this clustering approach. Despite the successful identification of protein families, only a few families covering sequences with fewer than 30 AAs could be identified (Fig. 6). This is due to several limitations, including the small number of collected sequences of this length in sORFdb (Fig. 2), too few small proteins reported in the literature being included in databases, and possible low sequence conservation [1]. Most families cover small proteins with a length between 40 to 50 amino acids and around 30 amino acids. This is likely due to the fact that there is an increased number of non-redundant small proteins of about 30 AA length in the sORFdb database which is caused by historical and technical length cutoffs (Fig. 2).

A functional description could be assigned to nearly all protein families, and HMMs with gathering cutoffs providing high accuracy were built accordingly. The various filtering steps employed during database creation and clustering reduced the number of false positives in the database and subsequently in the small protein families. For this reason the HMMs can be used to accurately predict small proteins for genome annotation. The most common functional descriptions of the small protein families were consistent with those reported in the literature, such as toxin-antitoxin systems, membrane-associated systems, and regulatory proteins [1, 12, 15, 18]. Despite this consistency of the identified high-quality small protein families, 37,024 small proteins were *a priori* excluded from the clustering by filtering strategies, and another 4,073 were reported as singletons. It is unclear whether these are true positives or false positives. They could be false positives that slipped through the extensive filtering steps during the database creation. This could be the case for pervasively translated non-functional small proteins predicted with ribosome profiling [21–23] or for false positives sORFs detected by gene prediction tools [5, 6]. Another possibility is that they may be small proteins without homologs in the protein databases or are underrepresented in bacterial genomes due to difficult detection.

sORFdb provides a comprehensive resource for information on practically all currently known high-quality sORFs and small proteins. Due to the understudied nature of these targets, there is still room for improvement in their detection and identification of their functions. Improved computational gene prediction and laboratory protocols for the identification of non-spurious small proteins and the elucidation of sORF and small protein properties are still open fields that need further research.

## 5 Conclusion

To the best of our knowledge, sORFdb is the first comprehensive, taxonomically independent database dedicated to sORF and small protein sequences and related information in bacteria. For this purpose, high-quality information from protein and genome databases enriched with physicochemical properties was combined. Furthermore, small protein families identified by a custom graph clustering approach accompanied by HMMs are provided to foster detection and consistent annotation.

In conclusion, the sORFdb database aims to serve as a high-quality primary resource for researchers studying sORFs and short proteins. It will help to improve the functional annotation of sORFs and small proteins, as well as the future detection of novel short proteins in bacteria.

## List of abbreviations

AA: Amino acid

HMM: Hidden Markov model

RBS: Ribosomal binding site

sORF: Short open reading frame

SRV: Score ratio value

**Supplementary information.** The article has one supplementary file.

## Declarations

### Ethics approval and consent to participate

Not applicable

### Consent for publication

Not applicable

### Availability of data and materials

The website can be accessed at https://sorfdb.computational.bio/. All data is available for download at https://zenodo.org/records/10688271. The source code of the sORFdb workflow and the clustering approach is available at https://github.com/ag-computational-bio/sorfdb.

## Authors’ contributions

JH and OS designed the study. JH designed, implemented, and tested the database creation workflow and clustering approach. JH, FC and SD conceived and designed the clustering approach. JH and LJ developed the website. LJ developed the server. JH conducted data analyses and interpreted the data. JH wrote the manuscript. AG, OS and SD substantially revised the manuscript. AG was responsible for funding. All authors read and approved the manuscript.

## Acknowledgements

The authors would like to thank Dr. Jochen Blom for his helpful advice on the score ratio values.

