## Supplementals for "sORFdb – A database for sORFs, small proteins, and small protein families in bacteria"

small proteins, protein families, short open reading frames, sORF, database, bacteria

**URLs:**

GitHub: https://github.com/ag-computational-bio/sorfdb

Zenodo: DOI 10.5281/zenodo.10688271

Web: https://sorfdb.computational.bio/

### Supplemental Tables

##### Table 1: Versions of the tools used for the download of the databases

| Tool | Version |
|---|---|
| bioawk | 1.0 |
| curl | 8.2.1 |
| diamond | 2.1.8 |
| gawk | 5.1.0 |
| grep | 3.11 |
| hmmer | 3.3.2 |
| lxml | 4.9.3 |
| pigz | 2.6 |
| python | 3.11.4 |
| rename | 1.601 |
| tar | 1.34 |
| wget | 1.20.3 |
| xopen | 1.7.0 |

##### Table 2: Versions of the tools used for the processing of the databases

| Tool | Version |
|---|---|
| biopython | 1.81 |
| diamond | 2.1.8 |
| ijson | 3.2.3 |
| jq | 1.6 |
| krona | 2.8.1 |
| peptides | 0.3.1 |
| polars | 0.16.14* |
| pyhmmer | 0.9.0 |
| pyrodigal | 2.1.0 |
| python | 3.10.12 |
| xopen | 1.7.0 |

*The python script taxonomy2krona.py was executed with version 0.19.0.

**Table 3: Versions of the tools used for the clustering**

| Tool | Version |
|---|---|
| blast | 2.14.1 |
| mcl | 22.282 |
| muscle | 5.1 |
| numpy | 1.25.2 |
| pandas | 2.0.3 |
| polars | 0.18.15 |
| pyhmmer | 0.10.2 |
| python | 3.10.12 |
| xopen | 1.7.0 |

**Table 4: Top 20 non-redundant small protein product annotations without ribosomal proteins**

| Product | Count |
|---|---|
| helix-turn-helix domain-containing protein | 86298 |
| helix-turn-helix transcriptional regulator | 73168 |
| transcriptional regulator | 66725 |
| acyl carrier protein | 66706 |
| antitoxin | 61763 |
| transposase | 51557 |
| type ii toxin-antitoxin system rele/pare family toxin | 41659 |
| dna-binding protein | 35427 |
| cold-shock protein | 33735 |
| sec-independent protein translocase protein tata | 31886 |
| xre family transcriptional regulator | 28640 |
| ferredoxin | 26492 |
| acylphosphatase | 26294 |
| hu family dna-binding protein | 26157 |
| type ii toxin-antitoxin system hica family toxin | 25448 |
| translation initiation factor if-1 | 25091 |
| co-chaperonin groes | 24978 |
| exodeoxyribonuclease 7 small subunit | 24499 |
| exodeoxyribonuclease vii small subunit | 24455 |
| type ii toxin-antitoxin system phd/yefm family antitoxin | 24142 |

#### Supplemental Figures

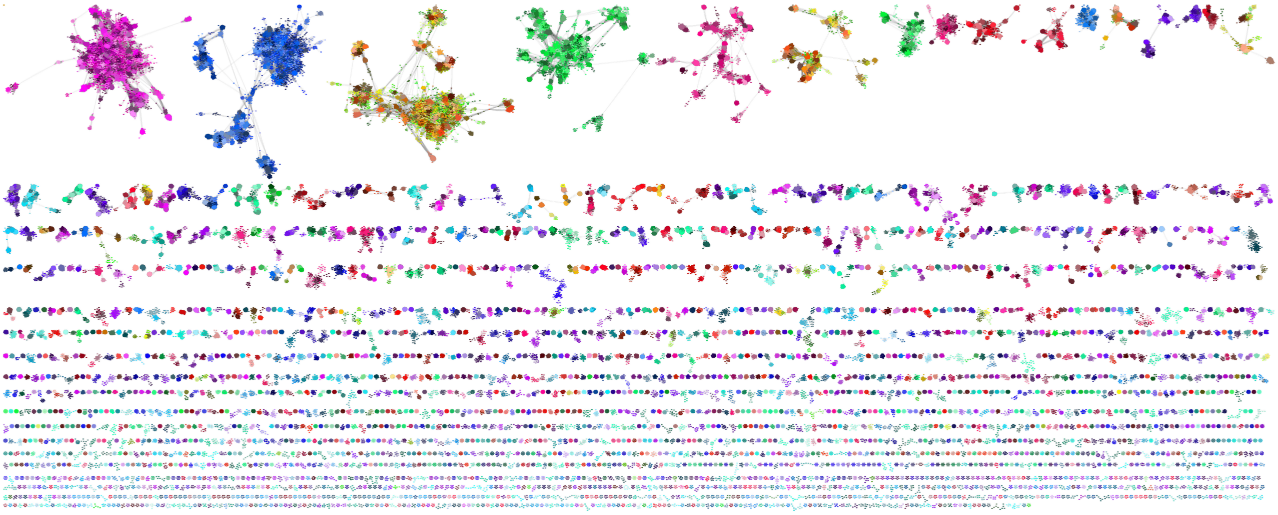

**Figure 1: Visualization of the graph network of identified small protein families**

All unconnected components of the graph were assigned a color palette. For the small protein families, one color is assigned in each component, where colors can occur multiple times. Families of one color separated by families of different colors are independent from another.
